# Liraglutide protects against diastolic dysfunction and improves ventricular protein translation

**DOI:** 10.1101/2022.08.23.504997

**Authors:** Cody Rutledge, Angela Enriquez, Kevin Redding, Mabel Lopez, Steven Mullett, Stacy Wendell, Michael Jurczak, Eric Goetzman, Brett A. Kaufman

## Abstract

**Purpose:** Diastolic dysfunction is an increasingly common cardiac pathology linked to heart failure with preserved ejection fraction. Previous studies have implicated glucagon-like peptide 1 (GLP-1) receptor agonists as potential therapies for improving diastolic dysfunction. In this study, we investigate the physiologic and metabolic changes in a mouse model of angiotensin II (AngII) mediated diastolic dysfunction with and without the GLP-1 receptor agonist liraglutide (Lira).

**Methods:** Mice were divided into sham, AngII, or AngII + Lira therapy for 4 weeks. Mice were monitored for cardiac function, weight change, and blood pressure at baseline and after 4 weeks of treatment. After 4 weeks of treatment, tissue was collected for histology, protein analysis, targeted metabolomics, and protein synthesis assays.

**Results:** AngII treatment causes diastolic dysfunction when compared to sham mice. Lira partially prevents this dysfunction. The improvement in function in Lira mice is associated with dramatic changes in amino acid accumulation in the heart. Lira mice also have improved markers of protein translation by Western blot and increased protein synthesis by puromycin assay, suggesting that increased protein turnover protects against fibrotic remodeling and diastolic dysfunction seen in the AngII cohort. Lira mice also lost lean muscle mass compared to the AngII cohort, raising concerns about peripheral muscle scavenging as a source of the increased amino acids in the heart.

**Conclusions:** Lira therapy protects against AngII-mediated diastolic dysfunction, at least in part by promoting amino acid uptake and protein turnover in the heart. Liraglutide therapy is associated with loss of mean muscle mass, and long-term studies are warranted to investigate sarcopenia and frailty with liraglutide therapy in the setting of diastolic disease.

## Introduction

Heart failure with preserved ejection fraction (HFpEF) is an increasingly prevalent syndrome in the United States, comprising over half of all heart failure diagnoses (1). While traditional heart failure therapies may help manage the symptoms of HfpEF, no pharmaceutical interventions have been shown to lower mortality in HfpEF patients (2). HfpEF is typified by signs and symptoms of fluid overload caused by impaired left ventricular relaxation, called diastolic dysfunction, and is associated with cardiac hypertrophy and fibrosis (3). Mouse studies of hypertrophy (4) and HfpEF-related models (5,6) have found that treatment with glucagon-like peptide 1 (GLP-1) agonists may improve markers of diastolic dysfunction. The mechanism of this improvement, however, remains unclear. Previous works have implicated changes to glucose regulation, cardiac bio-energetics, reduced inflammation, and altered endothelial function as possible mediators of GLP-1 agonist benefits in cardiovascular disease.(7,8). In this manuscript, we demonstrate that GLP-1 agonists alter proteostasis to improve cardiac function in diastolic disease.

GLP-1 is an incretin hormone released in the intestine that stimulates insulin secretion and lowers blood sugar (9,10). GLP-1 receptor agonists, such as liraglutide, improve outcomes in patients with type 2 diabetes mellitus by normalizing blood glucose levels (9,11). Additionally, GLP-1 agonists are known to improve cardiac outcomes in diabetic patients, demonstrating reduced major adverse cardiac events across multiple GLP-1 analogs (12,13). While liraglutide did not improve outcomes in heart failure with reduced ejection fraction patients in the FIGHT (14) and LIVE (15) trials, the effects on HFpEF remain less clear, and long term studies of the effect of GLP-1 agonists in HFpEF are on-going(16). GLP-1 receptor agonists have also been shown to improve cardiac outcomes in mouse models of hypertension (17), hypertrophy (4), HFpEF (5,6), and myocardial infarction (18). Additionally, GLP-1 receptor agonists have been approved for weight loss independent of diabetic status, and their use is becoming increasingly popular in non-diabetic patients (19–21). Given the broad application of GLP-1 agonists and the multiple cardiac effects this class of drugs exhibits, understanding the metabolic effects of GLP-1 agonists and possible benefits toward diastolic dysfunction is warranted.

This study investigated the cardiac metabolic changes in a mouse model with diastolic dysfunction and how that changed with GLP-1 therapy. We implanted mice with angiotensin-II (AngII) pumps for 4 weeks to model diastolic dysfunction, as previously described (22,23). A cohort of AngII mice was treated simultaneously with the GLP-1 receptor agonist liraglutide (Lira). We evaluated these mice for markers of diastolic dysfunction and profiled metabolomic changes to understand better the mechanism underlying GLP-1 targeted therapy in the heart.

## Methods

### Angiotensin Pump Implantation and Liraglutide Therapy

All mice were bred and housed at the University of Pittsburgh according to Institution Animal Care and Use Committee (IACUC; 21038951) protocols. Mice were kept on a 12-hour light/dark cycle with water and standard chow ad libitum. Eight-week-old male C57BL/6J mice were anesthetized using vaporized isoflurane (3%) and then underwent osmotic pump (Alzet, Cupertino CA) implantation or sham procedure. Pumps were pre-loaded with angiotensin released at a rate of 1.1 mg/kg/day. Pumps were weighed before and after loading to ensure complete filling and submerged overnight in warm saline before implantation. Mice were anesthetized, and the hair over the mid-back was removed using electrical clippers and sterilized with betadine; then, a skin incision was made in the implantation mice. Using forceps, a small subcutaneous pocket was opened, and an AngII-loaded pump was inserted. The incision was closed with a 5-0 vicryl dissolvable suture and covered with topical antibiotic ointment. Animals were monitored post-operatively for recovery for two hours and then daily after surgery. Sham mice were anesthetized but received no surgical incision or pump placement. The AngII mice were randomly assigned to either no treatment or liraglutide (Lira) groups. The Lira group received daily subcutaneous injections at 0.2 mg/kg/day. Lira animals were treated for a total of 4 weeks. No mice were excluded from the study.

### Echocardiography

Mice were anesthetized using isoflurane (1-3%) delivered by nose cone at baseline (8 weeks old) and four weeks after pump implantation. Transthoracic echocardiography was performed using a Vevo 3100 imaging system (Visual Sonics, Toronto, Canada) with a 40 MHz probe, as we previously described (24). Briefly, animals were maintained at 37 °C by a heating pad and were electrocardiogram (ECG) monitored using surface limb electrodes. Heart rate was maintained between 400-500 bpm during imaging by adjusting isoflurane concentration. M-mode and B-mode images of the heart were obtained for at least ten cardiac cycles along the parasternal long axis and left ventricular (LV) short axis. M-mode and Doppler imaging were obtained at the apical window. LV end-diastolic diameter, LV end-systolic diameter, and septal and free wall (systolic and diastolic) thickness were measured along the short axis. LV ejection fraction, LV fractional shortening, and corrected LV mass were calculated based on these values according to previously published Vevo-sonic equations (25). Transmitral Doppler imaging was used to calculate E and A waves as well as isovolumic contraction time (IVCT), isovolumic relaxation time (IVRT), and ejection time (ET). Tissue Doppler from the mitral annulus measured E’ and A’ values. Speckle tracking was performed by analyzing epicardial and endocardial tissue along the parasternal long-axis using Vevo 3100 VevoStrain software (26). The mean global longitudinal strain rate and reverse peak longitudinal strain rate were semi-automatically calculated during LV filling, as previously reported (27). Image analysis was performed by a blinded sonographer.

### Blood Pressure

Blood pressure was measured in conscious mice using a non-invasive CODA tail-cuff system (Kent Scientific, Torrington, CT). Mice were placed in a plastic holder and allowed to rest for 5 minutes before undergoing 5 acclimatization cycles, followed by 15 blood pressure measurement cycles. Systolic, diastolic, and mean blood pressure were calculated by the CODA software system. Animals were only included in the analysis if the system registered >50% of blood pressure readings as successful.

### Glucose Tolerance Tests and Body Composition

At 12 weeks of age, animals from all groups underwent a glucose tolerance test and body composition analysis. Mice were fasted for 6 hours before measuring baseline glucose concentrations in whole blood via the tail vein as previously described (28). D-glucose diluted in saline was then administered at 1 g/kg body weight by intraperitoneal (IP) injection. Blood glucose concentrations were measured every 15 min for 2 hours using Contour blood glucose test strips (Ascensia Diabetes Care US Inc, Parsippany, NJ). Body composition analyses were completed using ^1^H magnetic resonance spectroscopy (EchoMRI-100H, Houston, TX) after 4 weeks of treatment to compare changes in lean muscle mass, free water, and body fat in sham, AngII, and Lira groups.

### Tissue and Serum Collection

Following the four-week echocardiography, mice were euthanized under isoflurane anesthetic. A cardiac puncture was performed for blood collection using a heparinized syringe and centrifuged for plasma isolation, which was then flash-frozen. The heart was rapidly excised and washed with chilled saline, then dissected into the atria, right ventricle, and left ventricle before flash freezing in liquid nitrogen or fixing in formalin for tissue histology. Gastrocnemius muscles were collected and snap-frozen at the time of sacrifice.

### Tissue Histology

Hearts were fixed overnight in 10% formalin (Thermo, #SF100) at 4°C, then washed with PBS and transferred into 70% ethanol at room temperature to dehydrate the sample. Tissues were embedded in paraffin before sectioning at 4 µm by the Histology Core at the Children’s Hospital of Pittsburgh. Sections were stained with Masson’s trichrome. Histological evaluation was performed in a blinded fashion at 40x magnification. Images were obtained on a Nikon Eclipse E800 (Nikon, Tokyo, Japan) microscope at 40x magnification. Images were captured via Nikon NIS-elements software and analyzed using ImageJ. To investigate cardiomyocyte size, 50 short-axis cardiomyocytes were measured for diameter.

### Metabolomics Analysis

Targeted metabolomic studies were completed for all three groups via liquid chromatography-high resolution mass spectrometry (LC-HRMS) for citric acid cycle and glycolytic metabolites, amino acids, carnitines, and acetyl-CoA. Metabolic quenching and polar metabolite extraction were performed using ice-cold 80% methanol/0.1% formic acid at 500 µL per 50 mg of frozen, powdered LV tissue. An internal standard mix containing (D3)-creatinine and (D3)-alanine, (D4)-taurine, and (D3)-lactate (Sigma-Aldrich) was added to the sample lysates for a final concentration of 100 µM. After 3 min of vortexing, the supernatant was cleared of protein by centrifugation at 16,000 × *g*. Cleared supernatant (3 µL) was subjected to online separation and analysis by LC-HRMS. In brief, samples were injected via a Thermo Vanquish UHPLC and separated over a reversed-phase Thermo HyperCarb porous graphite column (2.1 × 100 mm, 3-μm particle size) maintained at 55°C. For the 20-min LC gradient, the mobile phase consisted of solvent A (water/0.1% formic acid) and solvent B (acetonitrile/0.1% formic acid). The gradient was the following: 0–1 min 1% B, increasing to 15% B over 5 min, increasing to 98% B over 5 min, and held at 98% B for 5 min before equilibration to starting conditions. The Thermo ID-X tribrid mass spectrometer was operated in positive and negative ion mode, scanning in full MS mode (two microscans) from 100 to 800 *m/z* at 70,000 resolution with an automatic gain control target of 2 × 10^5^. The source ionization setting was 3.0 kV and 2.4vkV spray voltage for positive and negative modes, respectively. Source gas parameters were 45 sheath gas, 12 auxiliary gas at 320°C, and 8 sweep gas. Calibration was performed before analysis using the Pierce FlexMix Ion Calibration Solution (Thermo). Alignment and peak area integration were then extracted manually using Quan Browser (Xcalibur ver. 2.7; Thermo Fisher). Atomic percent enrichment was calculated using the established mass isotopomer multiordinate spectral analysis (MIMOSA) method to remove the natural [^13^C] abundance background. Data are graphically represented using Clustvis-generated heatmaps (29). Raw data were uploaded to the UCSC Metabolomics Workbench as supported by the Common Fund of the National Institutes of Health (Track ID 3485).

### Western Blot, Oxyblot, and Serum ELISA

Frozen LV tissue was homogenized in lysis buffer containing a protease/phosphatase cocktail (Sigma-Aldrich, St. Louis, MO, #11697498001) and normalized for protein content using a BCA assay (Life Technologies, Carlsbad, CA, #23235). Samples were separated on NuPage 4-12% gradient SDS-PAGE gels (ThermoFisher, Waltham, MA, #WG1403BOX) and transferred onto iBlot nitrocellulose membranes (Invitrogen, #IB301001). Membranes were blocked in 5% milk (non-phosphorylated antibodies) or 5% BSA (phosphorylated antibodies) for 1 hour and then incubated overnight at 4°C with primary antibodies p-S6 (Ser240/244, 1:1000, Cell Signaling, #2215), S6 (1:1000, Cell Signaling, #2217), GAPDH (1:5000, Millipore, St. Louis, MO, #AB2302), p-4EBP1 (Thr37/46, 1:1000, Cell Signaling, #2855), 4EBP1 (1:1000, Cell Signaling, #9644), P62 (Invitrogen #PA5-20839, 1:1000), Ubiquitin (Invitrogen #1301600, 1:1000), CPT1b (Thermo #PA5-79065, 1:1000), MCAD and VLCAD (both courtesy of Vockley laboratory, 1:1000), phospho-BCKDH (Cell Signaling #40368, 1:1000) and BCKDH (Cell Signaling #90198, 1:1000). Following incubation, membranes were washed with TBS-tween and then probed for 1 hour at room temperature with anti-mouse or anti-rabbit secondary antibodies (Jackson ImmunoResearch, West Grove, PA, #115-035-003 and #115-035-144). Oxyblot (Millipore #S7150) detection of carbonyl groups was performed per manufacturer recommendations. All images were obtained on a ChemiDoc XRS imaging system (BioRad, Hercules, CA) and analyzed using ImageJ software (National Institutes of Health, Bethesda, MD). Plasma was evaluated for BNP using a Mouse NT-proBNP ELISA kit (Novus Biologicals, Centennial, CO, NBP2-70011) and read on a plate spectrometer.

### Puromycin Assay for Protein Synthesis

Thirty minutes before animal sacrifice, a cohort of mice from each treatment group was given an IP injection of puromycin (40 nm/g) to assess protein synthesis in the heart (30,31). Hearts were flash frozen and later homogenized for Western blot analysis as described above. The transfer membrane was blocked with milk and then incubated overnight at 1:1000 of PMY-2A4 primary antibody (Developmental Studies Hybridoma Bank, University of Iowa), followed by 1 hour with anti-rabbit secondary and developed. Whole lane densitometry was analyzed and compared between groups.

### Fatty Acid Oxidation (FAO)

Fresh LVs were homogenized in BIOPS buffer (10 mM Ca-EGTA, 0.1 µM calcium, 20 mM imidazole, 20 mM taurine, 50 mM K-MES, 0.5 mM DTT, 6.56 mM MgCl2, 5.77 mM ATP, 15 mM phosphocreatine to pH 7.1) and the lysate centrifuged at 500 g for 5 min to remove debris. FAO reactions consisted of 5 uL of lysate in a total reaction volume of 200 uL containing 200 mM sucrose, 10 mM Tris-HCl, 5 mM KH_2_PO_4_, 0.2 mM EDTA, 0.5 % fatty acid-free BSA, 80 mM KCl, 1 mM MgCl_2_, 0.2 mM L-carnitine, 0.1 mM malate, 0.05 mM coenzyme A, 2 mM ATP, 1 mM DTT, and 125 uM of ^14^C-labeled palmitic acid (PerkinElmer). ^14^C-labeled products were separated from the reactions by extraction with chloroform/methanol as previously described (32). The rate of fatty acid oxidation (FAO) was normalized to protein concentration.

### Statistical Analysis

Data are expressed as mean ± standard error in all figures. A p-value of ≤ 0.05 was considered significant for all comparisons. One-way ANOVA with Tukey’s multiple comparisons test was used to compare groups. All statistical analysis was completed using GraphPad Prism 8 software (San Diego, CA).

## Results

### AngII-treated mice develop diastolic dysfunction that can be prevented by Lira treatment

The angiotensin-aldosterone system activation directly contributes to diastolic dysfunction in humans (33). The effects of continuous AngII infusion has been shown to induce diastolic dysfunction (34)with some models progressing to hypertrophy and reduced contractility (35,36). As such, we needed to confirm that the effect of 1.1 mg/kg/day AngII-delivery was limited to diastolic dysfunction. Following four weeks of infusion, AngII-treated mice indeed developed diastolic dysfunction as measured by Doppler flow analysis of the mitral valve, including decreased E/A and E/E’ (Figure 1A). AngII mice also have reduced peak reverse longitudinal strain rate (rLSR) by speckle tracing analysis. These changes were all prevented in the AngII mice treated with Lira. AngII+Lira mice had unchanged E/A, E/E’, and rLSR compared to the sham mice but significantly improved E/A, E/E’, and rLSR compared to the untreated AngII cohort. There were no significant changes to isovolumic relaxation time (IVRT) between treatment groups. Ejection fraction (EF) was measured as an indicator of systolic function. There was no change in EF between treatment groups (Figure 1B).

**Figure 1.**
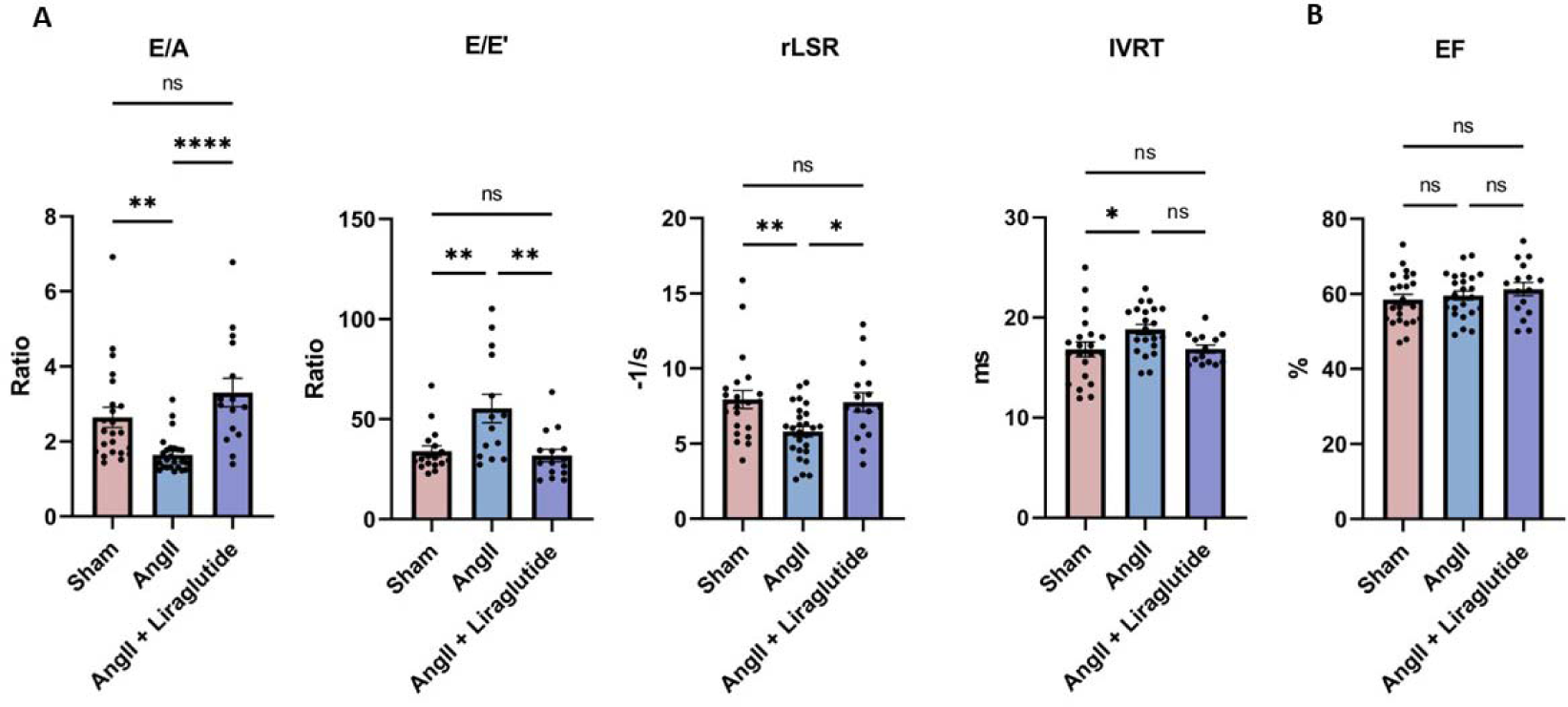
Liraglutide protects against angiotensin (AngII) induced diastolic dysfunction. A) AngII mice (n=14) have evidence of diastolic dysfunction when compared to sham (n=17) as measured by reduced E/A and elevated E/E’ by doppler imaging and decreased peak reverse longitudinal strain rate (rLSR) by speckle tracing. E/A, E/E’, and rLSR are all unchanged from sham in the AngII mice treated with liraglutide (n=15). Isovolumic relaxation time (IVRT) prolonged in the AngII mice compared to sham but not significantly changed in the AngII mice treated with liraglutide. B) Ejection fraction (EF) is unchanged between groups. *p ≤ 0.05, **p ≤ 0.01, ****p ≤ 0.0001 by ANOVA with Tukey’s multiple comparison test.

### AngII causes cardiac fibrosis that is partially prevented by Lira treatment

Diastolic dysfunction involves cardiac fibrosis, which contributes to the diminished relaxation of the heart (37). To confirm this effect of AngII and determine whether Lira altered its development, we measured fibrotic area using Masson Trichrome Stain (Figure 2A). As expected, AngII significantly increased blue staining, which stains collagen, while Lira treatment reduced blue staining to an intermediate level by either percentage or relative cross-sectional area. We conclude that Lira is acting upstream of the fibrotic effects of chronic AngII exposure.

**Figure 2.**
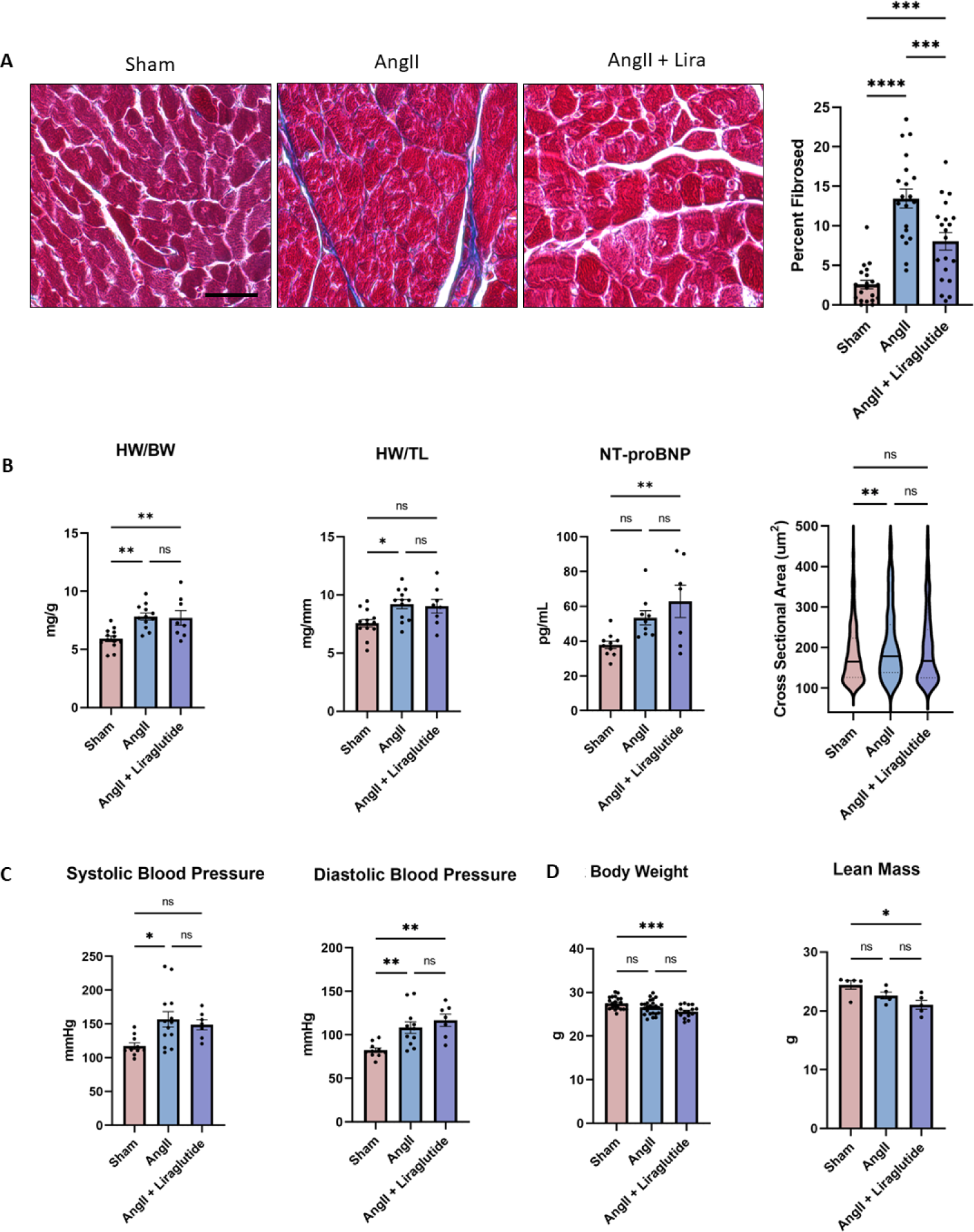
Changes to fibrosis, hypertrophy, blood pressure, and lean muscle mice. A) Left ventricles (LV’s) from AngII mice have increased fibrosis by trichrome staining when compared to sham mice. AngII+Liraglutide mice have less fibrosis when compared to AngII mice (n=20 high power fields from 4 mice per group). B) Both AngII and AngII+Liraglutide mice have elevated HW/BW than sham mice. HW/TL is elevated in AngII mice compared to sham, while serum NT-proBNP Is elevated in the AngII+Liraglutide mice compared to sham. AngII myocytes have increased cross-sectional area compared to sham that is unchanged in the AngII+Liraglutide mice (n=200-300 myocytes from 4 mice per group). C) Systolic blood pressure is elevated in AngII (n=12) compared to sham (n=10) but not significantly changed in AngII+Liraglutide mice (n=7). Diastolic blood pressure is significantly elevated in both AngII and AngII+Liraglutide mice compared to sham. D) Total body weight and lean muscle mass are decreased in AngII+Liraglutide mice when compared to sham (n=5 per group). *p ≤ 0.05, **p ≤ 0.01,***p ≤ 0.001 ****p<0.0001 by ANOVA with Tukey’s multiple comparison test. Scale bar = 75 um.

### Lira does not affect AngII-induced blood pressure elevation and hypertrophy

In pursuit of upstream mechanisms, we examined whether Lira modulated the blood pressure effects of AngII. As expected, we found that systolic and diastolic blood pressure (BP) is elevated in the AngII mice compared to sham (Figure 2B), comparable to previous models (22,35). While AngII+Lira mice had elevated diastolic BP compared to the sham group, there was no significant difference between sham and AngII+Lira systolic BP. AngII+Lira mice have unchanged systolic and diastolic BP compared to the untreated Ang mice. AngII mice developed hypertrophy, as noted by a significantly larger cardiomyocyte cross-sectional area in the AngII-treated mice than sham (Figure 2A). This result places the Lira protective mechanism downstream of the hypertensive trigger.

### AngII+Lira mice have lower body weight and lean mass than AngII mice

Elevated blood pressure is associated with body weight reduction in mice, regardless of whether the strain is prone to AngII-induced heart failure (38). Whether Lira might be acting on a body weight control mechanism. While animal body weights were unchanged at 8 weeks (Table 1), there was a significant decrease in 12-week body weight between the AngII+Lira mice and sham. While the AngII cohort did not have a significantly lower body weight at 12 weeks (4 weeks of AngII treatment), they did drop substantially in body weight over time compared to sham mice (Table 1). To quantify the body composition accounting for these changes, mice underwent whole-body MRI. There were no significant changes to body fat or free water composition between groups, but the AngII+Lira mice had lower total lean mass compared to sham (Figure 2C) that equaled the total body mass lost, suggesting that Lira treatment may be contributing to muscle loss in the challenged mice. The data suggests muscle loss may be the primary driver of weight loss in the Lira vs. sham mice.

**Table 1.**
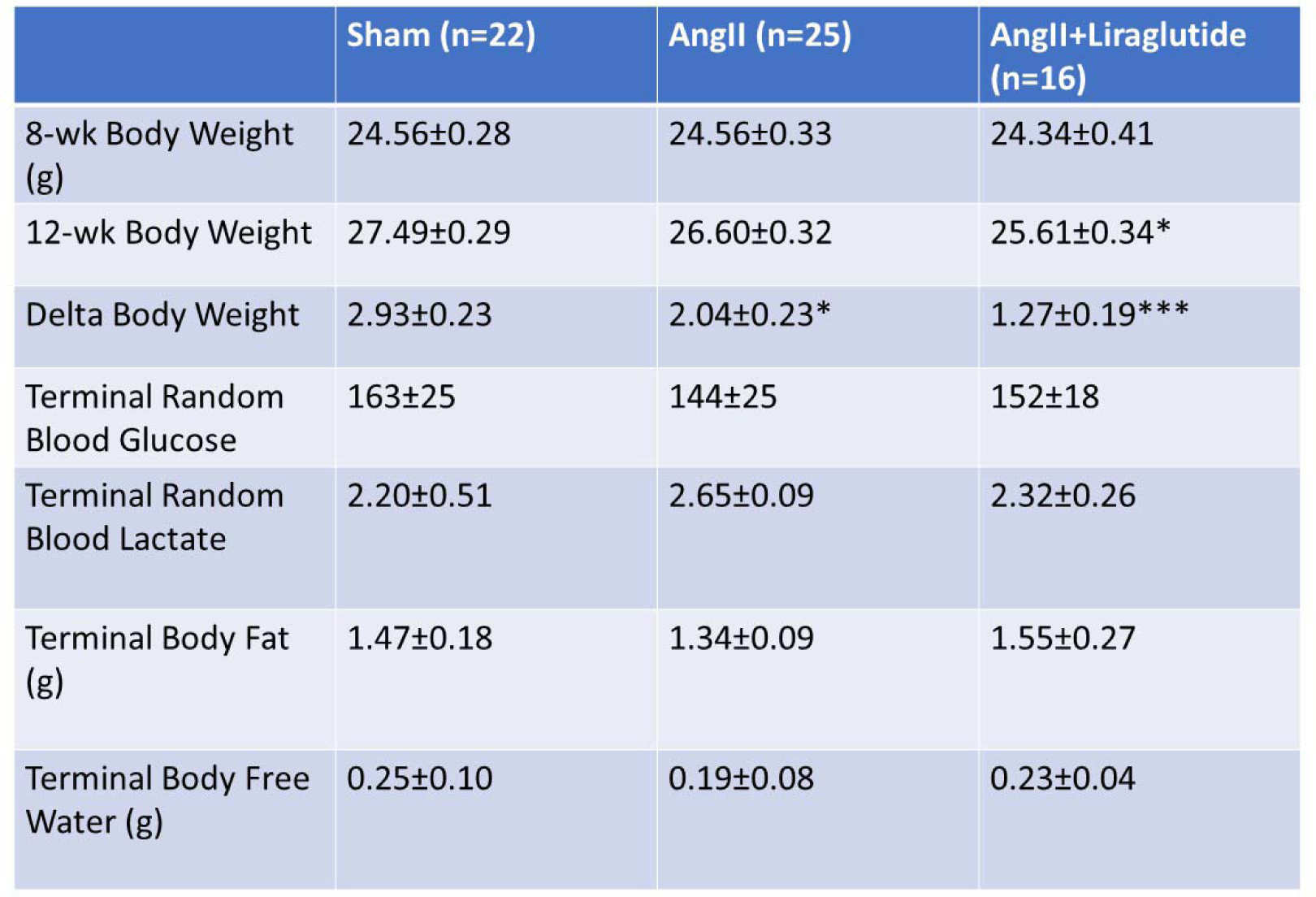
Physiologic characteristics of Sham, AngII, and AngII+Liraglutide mice. At the time of surgery (8-week-old mice), there is no change to baseline body weight between Sham (n=22), Ang (n=25), and AngII+liraglutide mice (n=16). 4-weeks after surgery, AngII+liraglutide mice have lower body weight than sham mice. Both AngII and AngII+Liraglutide have lower delta body weights over the 4-week experiment. There is no change to random blood glucose, lactate, body fat, or free water between groups at terminal procedure (n=5/group). *p ≤ 0.05, ***p ≤ 0.001 vs sham by ANOVA with Tukey’s multiple comparison test.

|  | Sham (n=22) | AngII (n=25) | AngII+Liraglutide (n=16) |
| --- | --- | --- | --- |
| 8-wk Body Weight (g) | 24.56±0.28 | 24.56±0.33 | 24.34±0.41 |
| 12-wk Body Weight | 27.49±0.29 | 26.60±0.32 | 25.61±0.34* |
| Delta Body Weight | 2.93±0.23 | 2.04±0.23* | 1.27±0.19*** |
| Terminal Random Blood Glucose | 163±25 | 144±25 | 152±18 |
| Terminal Random Blood Lactate | 2.20±0.51 | 2.65±0.09 | 2.32±0.26 |
| Terminal Body Fat (g) | 1.47±0.18 | 1.34±0.09 | 1.55±0.27 |
| Terminal Body Free Water (g) | 0.25±0.10 | 0.19±0.08 | 0.23±0.04 |

### AngII+Lira mouse hearts have altered amino acid accumulation compared to sham and AngII

Prior work showed that Lira treatment of *ex vivo* hearts showed no acute impact on glycolysis, glucose oxidation and fatty acid oxidation, while one-day systemic Lira treatment increased cardiac glucose oxidation in lean mice (7). However, precisely what impact Lira has on myocardial intermediate metabolism in the context of the development of diastolic dysfunction is unclear. To assess the metabolite changes related to AngII and AngII+Lira treatments, homogenized hearts underwent targeted metabolomic profiling for amino acids, citric acid cycle, glycolytic intermediates, ketones, and carnitines (Figure 3 and Supplemental Figure 2). AngII+Lira mice were found to have a dramatic increase in amino acid accumulation (Figure 3A). 23 of the 26 amino acids quantified were elevated in the AngII+Lira cohort compared to sham mice, and 17 of the 26 amino acids were elevated compared to AngII mice (Supplemental Figure 2). Interestingly, we note a much less dramatic amino acid accumulation in plasma samples of the AngII+Lira mice when compared to sham (6 of 26 amino acids significantly changed, Supplemental Figures 3 and 4) or AngII mice (1 of 23 amino acids), suggesting that the amino acid change is not specific to the myocardium, but is occurring to a lesser extent in circulating plasma as well.

**Figure 3.**
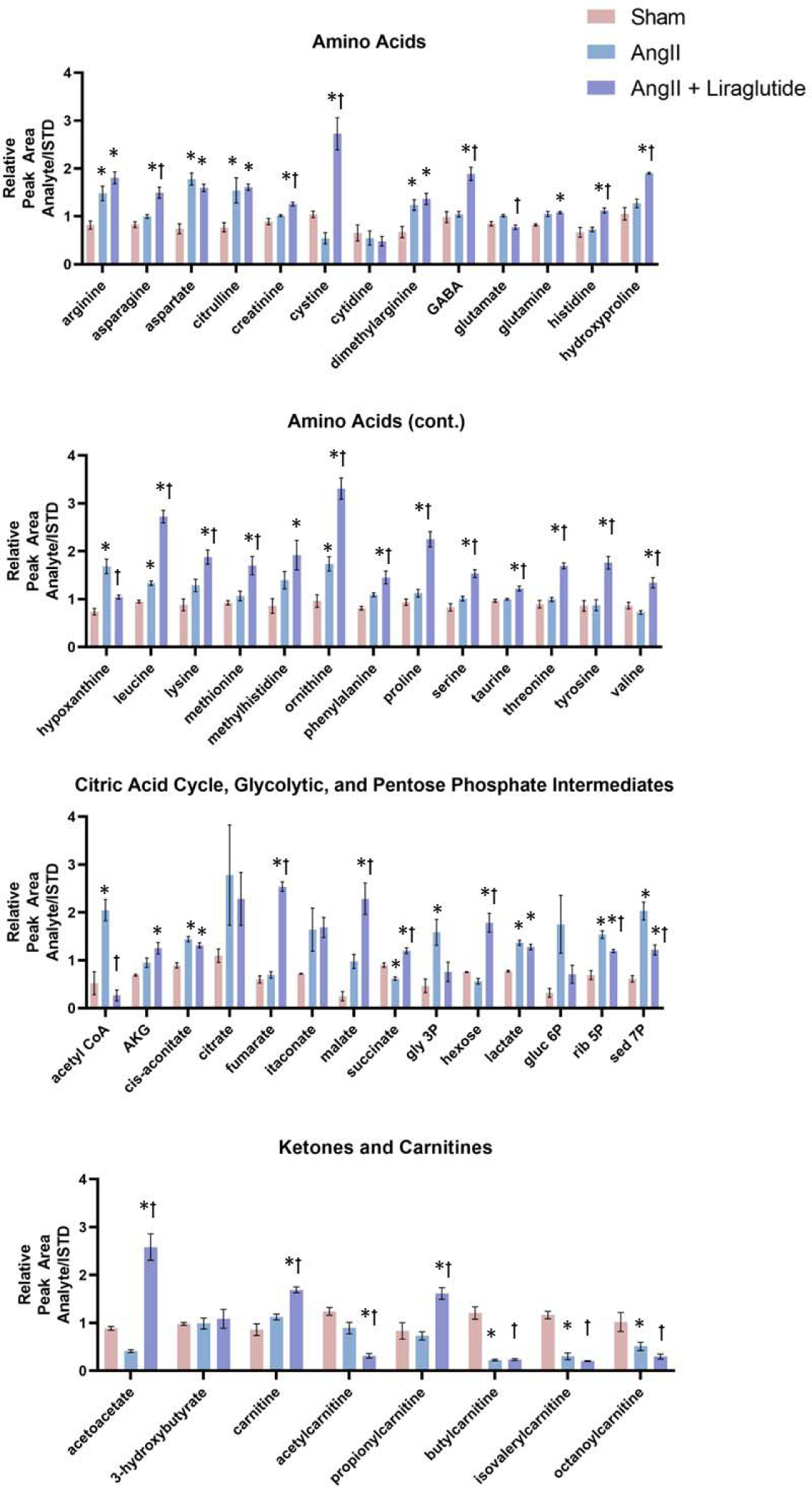
Targeted analysis of amino acids, ketones, citric acid cycle, glycolytic and pentose phosphate intermediates, and carnitines by metabolomic evaluation. Quantification of amino acids, citric acid cycle, glycolytic, and pentose phosphate intermediates, ketones in sham, AngII, and AngII+Lira mice (n=3-4 per group). *p ≤ 0.05 vs sham, ꝉp ≤ 0.05 vs AngII by ANOVA with Tukey’s multiple comparison test. gly 3p= glyceraldehyde-3-phosphate, rib 5p= ribulose-5-phosphate, gluc 6p= glucose 6 phosphate, sed 7p= sedoheptulose 7-phosphate.

### Citric acid cycle, glycolytic, and pentose phosphate intermediates are altered by AngII and AngII + Liraglutide treatments

Fewer changes among groups were noted when comparing the abundance of the citric acid cycle, glycolytic, and pentose phosphate intermediates in the LVs (Figure 3C-E). In particular, acetyl-CoA was significantly elevated in the AngII cohort compared to sham (Supplemental Figure 2) but normalized to sham levels in the AngII+Lira cohort. AngII mice have altered levels of cis-aconitate and succinate, but otherwise citric acid components tested were unchanged from sham. AngII+Lira hearts, however, have nearly all citric acid components elevated when compared to sham mice. In the glycolytic and pentose phosphate pathway metabolites, glyceraldehyde-3-phosphate, glucose-6-phosphate, ribulose-5-phosphate, and sedoheptulose 7-phosphate were elevated in AngII mice compared to sham. Relative to sham, the AngII + Lira mice have less dramatic changes than the AngII cohort but show elevated hexose, lactate, ribulose-5-phosphate and sedoheptulose 7-phosphate. These changes were much less pronounced in plasma metabolomics, as only cis-aconitate, citrate, itaconate, and ribulose-5-phosphate levels were significantly altered in the AngII mice compared to sham mice (Supplemental Figure 3 and 4).

### Ketone and carnitine profiles are altered between treatment groups

Increased ketone and free fatty acid acid update has been observed in heart failure (39) and has been suggested targets for HFpEF (40)), but hasn’t been looked at in a mouse model of diastolic dysfunction. For myocardial ketones, we found that acetoacetate levels were elevated in the AngII+Lira mice when compared to both sham and AngII mice (Figure 3 and Supplemental Figure 2), while 3-hydroxybutyrate levels were unchanged. For short and long chain acylcarnitines, we found that carnitine and propionylcarnitine levels were elevated in the AngII+Lira compared to sham, while acetylcarnitine, butylcarnitine, isovalerylcarnitine, and octanylcarnitine levels were all significantly reduced between the same cohorts. In contrast, plasma analysis showed no significant changes between sham and AngII+Lira mice except for 3-hydroxybutyrate.

### AngII+Lira mouse hearts have altered markers of mRNA processing, autophagy, and protein synthesis

To better understand the role of the amino acid accumulation in the heart, we performed Western blot analysis for markers of mRNA translation and autophagy (Figure 4). We found that phosphorylated ribosomal protein S6, which positively correlates with ribosomal protein translation (41), is elevated in the AngII+Lira mice when compared to sham (Figure 4A). Similarly, we quantified the translation repressor 4EBP1. When 4EBP1 is phosphorylated, it is degraded, allowing eIF4E to initiate translation (42). p-4EBP1/4EBP1 is elevated in the AngII mice compared to sham; however, it has significantly higher expression in AngII+Lira mice compared to the AngII group (Figure 4A). p62, a receptor protein associated with the degradation of ubiquitinated proteins and autophagy (43), is elevated in AngII+Lira mice when compared to both sham and AngII groups (Figure 4B). To assess protein synthesis, we treated a cohort of mice with the antibiotic puromycin, which incorporates into elongating amino acid chains and can be quantified as a marker of protein synthesis (30). We found that AngII mice had higher puromycin incorporation than sham mice (Figure 4C). AngII+Lira mice had significantly higher puromycin incorporation than both the sham and AngII cohorts, indicating the rate of translation was highest in this group. Finally, to assess markers of protein degradation and oxidative damage, we performed Western blot analysis of total ubiquitination and carbonylation by Oxyblot, respectively. We found that ubiquitination is elevated in both AngII and AngII+Lira mice compared to sham (Supplemental Figure 5A) and that carbonylation is elevated in AngII mice compared to sham, but unchanged in the AngII+Lira cohort (Figure 5B), which may suggest increased protein damage and degradation in both the AngII and AngII+Lira cohorts compared to sham.

**Figure 4.**
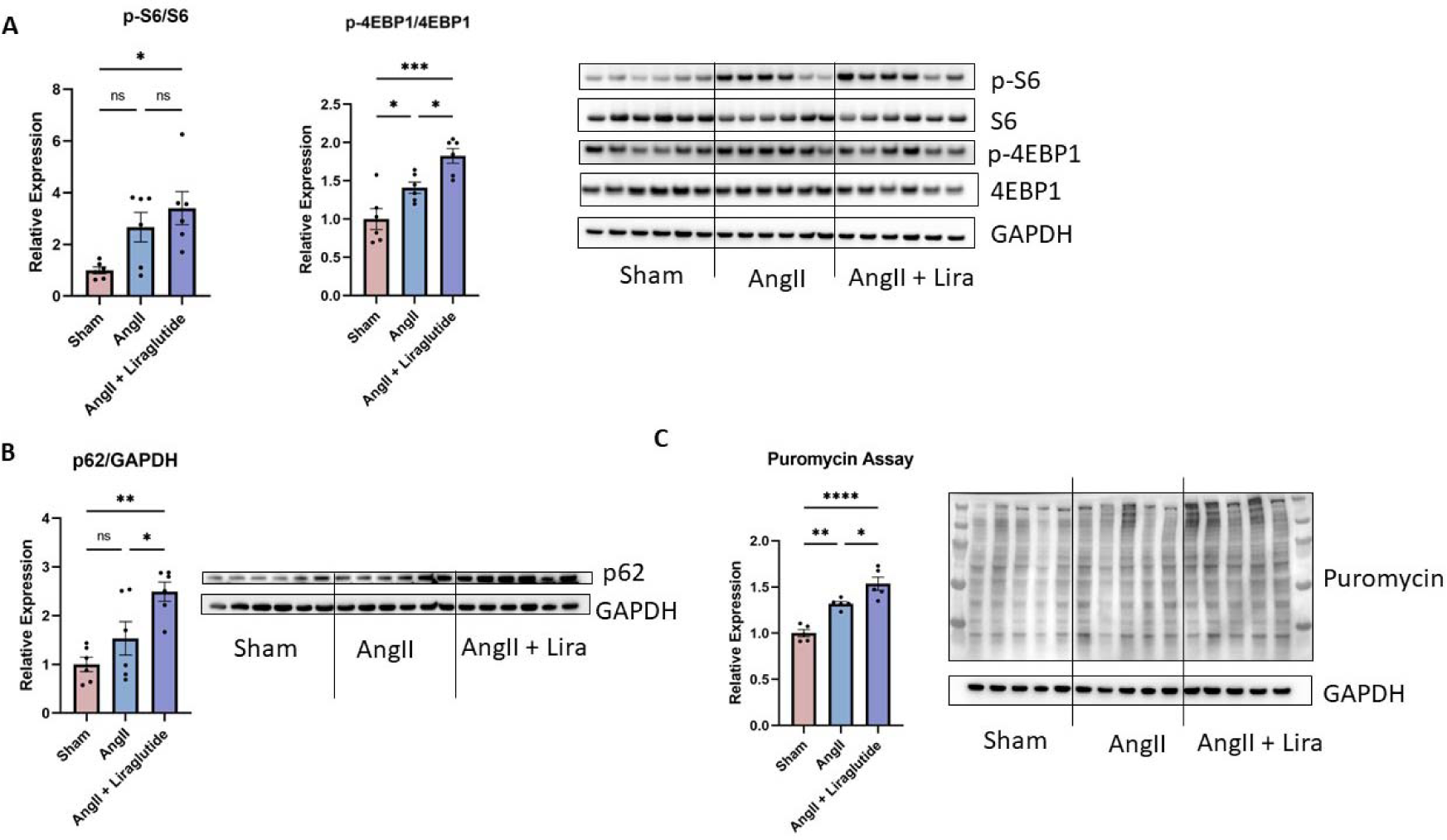
Liraglutide increases markers of mRNA translation and autophagy and promotes protein synthesis in AngII mice. A) AngII mice treated with liraglutide have elevated p-S6/S6 compared to sham mice. AngII mice treated with liraglutide have elevated p-4EBP1/4EBP1 compared to both sham and AngII mice (n=6/group) B) AngII mice treated with liraglutide have elevated p62 compared to Sham and AngII mice (n=6/group). C) Puromycin inclusion is increased in AngII mice treated with liraglutide (n=5/group) when compared to Sham and AngII mice. *p ≤ 0.05, **p ≤ 0.01,***p ≤ 0.001 ****p ≤ 0.0001 by ANOVA with Tukey’s multiple comparison test.

**Figure 5.**
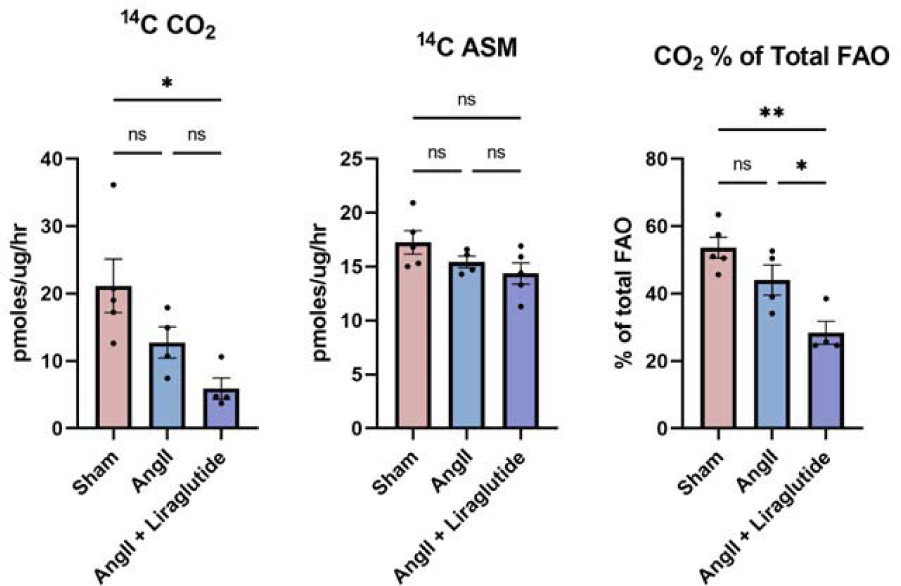
Palmitate oxidation is altered in AngII and AngII + Liraglutide mice. ^14^C labeled palmitate was given to homogenized hearts in each group to measure fatty acid oxidation (FAO). AngII + Liraglutide mice had significantly less CO2 formation than sham mice. There was no change to acid soluble metabolites (ASM) between groups. CO2 production was significantly lower in the AngII + Liraglutide mice when compared to Sham and AngII when comparing CO2 as a percentage of total palmitate oxidation. n=4/group, *p ≤ 0.05, **p ≤ 0.01 by ANOVA with Tukey’s multiple comparison test.

### Fatty Acid Utilization is reduced in AngII+Lira Mice

Given prior reports of genetically enhanced FAO preserves cardiac function in diastolic dysfunction (22), as well as the changes to carnitine profiles observed in our AngII and AngII+Lira hearts (Figure 3), we assessed FAO in all three groups using radiolabeled palmitate in homogenized hearts (Figure 5). Surprisingly, we found that AngII+Lira hearts had significantly reduced palmitate oxidation to CO_2_ compared to sham mice (Figure 5). No changes were observed for acid soluble metabolites (ASM), which include predominately acetyl-CoA and some citric acid cycle intermediates, between groups. By quantifying CO_2_ production compared to total FAO (sum of CO_2_ and ASM), we found that AngII+Lira mice had lower palmitate oxidation than sham or AngII mice. These data suggest that increased FAO is not the driver of improved cardiac function in AngII+Lira mice. Interestingly, we found no change to FAO enzymes by Western blot analysis, including medium-chain acyl-CoA dehydrogenase (MCAD), very long-chain acyl CoA dehydrogenase (VLCAD), or carnitine palmitoyltransferase 1B (CPT1B; Supplemental Figure 5), suggesting that the FAO changes are not driven by enzyme expression.

### Branched-chain amino acid (BCAA) utilization may be altered in AngII+Lira mice

Given the accumulation of branched-chain amino acids leucine and valine in the AngII+Lira mice (Figure 3A), we assessed for expression of branched-chain α-keto acid dehydrogenase complex (BCKDH), which is the rate-limiting step in BCAA degradation (44). We found that total BCKDH is decreased in AngII compared to sham but is restored in the AngII+Lira mice (Supplemental Figure 5D). Phosphorylated BCKDH (serine 293), which inactivates BCKDH (45), is not significantly changed in the AngII mice. However, p-BCKDH is elevated in the AngII+Lira cohort compared to both sham and AngII mice, suggesting reduced BCAA utilization.

## Discussion

AngII infusion is a well-studied mouse model of hypertension that has been associated with a variety of cardiac pathologies, ranging from subtle diastolic dysfunction to systolic heart failure (34,35,46). Here we show that our AngII treatment causes the development of diastolic dysfunction as determined by multiple echocardiographic modalities with no change to cardiac ejection fraction (Figure 1). These mice have elevated blood pressure, cardiac hypertrophy, and fibrosis. We found that daily treatment with liraglutide completely prevented diastolic dysfunction caused by the AngII treatment, with no significant changes to blood pressure or hypertrophy (Figure 2). A previous murine model treated with a slightly higher dose of AngII showed that liraglutide treatment prevented cardiac hypertrophy and systolic dysfunction, but diastolic parameters were not evaluated (4). That study did find less fibrosis in AngII+Lira compared to AngII, consistent with our results. Additional work also found that liraglutide treatment improves diastolic dysfunction and pulmonary congestion in a two-hit mouse model of HFpEF utilizing AngII treatment combined with a high-fat diet (6), and another has shown that liraglutide can prevent cardiomyocyte death in a pressure overload model of hypertrophy (47). All of these works note reduced cardiac fibrosis in liraglutide-treated hearts. Our work supports these findings that liraglutide functionally protects against AngII-mediated remodeling and fibrosis in the heart. However, whether this is a metabolic restoration or a bypass has not been investigated previously.

A novel finding in our study involves the metabolite changes observed in the AngII+Lira cohort. Targeted metabolomic studies of the heart found a dramatic accumulation of nearly all amino acids studied. AngII-treated mice had a nearly identical amino acid profile compared to the sham group, with the amino acid ornithine as the only significantly changed metabolite. AngII+Lira mice had 23 of 26 amino acids significantly elevated compared to the sham mice and 17 of 26 compared to the AngII cohort. We initially considered that this broad elevation of amino acids represented an increased reliance on amino acids as an energy substrate, particularly given the recent interest in branch-chain amino acids (BCAA) as a fuel source in cardiomyopathy and heart failure (48,49). This is supported by the significant changes to BCAA’s leucine and valine in our AngII+Lira mice; however, we show that p-BCKDH, which inactivates the rate-limiting step in BCAA catabolism, is elevated in the AngII+Lira cohort and argues against BCAA utilization as a fuel source in this cohort (Supplemental Figure 5). However, recent work on BCAA’s in the heart linked BCAA accumulation to increased protein synthesis and hypertrophy (50), leading us to investigate markers of protein synthesis in our model.

We found that phosphorylated S6, a marker of ribosomal activity (41), is elevated in the AngII+Lira mice compared to the sham group (Figure 4A). Similarly, AngII+Lira mice had increased phosphorylation of the translational repressor protein 4EBP1, which leads to the 4EBP1 degradation and increased mRNA translation (42). To measure translation directly, we injected mice with puromycin and found that protein synthesis is increased in the AngII+Lira compared to both sham and AngII cohorts. It should be noted that p-4EBP1 and puromycin incorporation were elevated in the AngII mice compared to sham, suggesting that AngII alone drives increased protein synthesis, potentially as a compensatory response to cardiac stress. Increased protein synthesis has been associated with pathologic cardiac hypertrophy (50). When treated with liraglutide, protein synthesis is further increased above AngII levels, potentially amplifying the compensatory effect and protecting against fibrosis and diastolic dysfunction. Previous work from our group has shown that increased cardiac protein synthesis is acutely protective in a model of ischemic damage (51,52). Despite the elevated protein synthesis in the AngII + Lira mice compared to the AngII mice alone, we did not see any worsened hypertrophy in the liraglutide cohort.

One possible explanation for the discrepancy between increased protein synthesis and unchanged hypertrophic measurements is that the liraglutide acts downstream of hypertrophy, allowing cardiomyocytes to persevere against the increased afterload related to elevated blood pressure. This concept is supported by elevated p62, a marker of autophagy, in the AngII+Lira mice (Figure 4B), as well as increased ubiquitination in the AngII and AngII+Lira mice compared to sham (Supplemental Figure 6A). Protein carbonylation, a marker of oxidative damage, is elevated in the AngII mice compared to sham but not significantly changed in the AngII+Lira group (Supplemental Figure 6B). Taken together, these data suggest that Lira is amplifying protein synthesis and turnover of damaged proteins to prevent fibrosis-induced diastolic dysfunction.

The increased protein synthesis and likely turnover observed in the AngII+Lira heart raises questions about the source of amino acids accumulating in the heart. All groups were fed *ad libitum* with a normal chow diet. Although food intake was not monitored in these mice, we found that the AngII+Lira mice had a reduced terminal weight compared to the sham group (Figure 2). Though this may be expected based on the role of GLP1-R agonists as a weight-loss agent (19,53), we did not find any significant changes to body fat or free water in the mice (Table 1). Rather, the decreased weight seems to be driven by a significant loss of lean muscle mass in the AngII+Lira cohort (Figure 2). GLP-1R agonists, particularly liraglutide, have been linked to reduced lean body mass in human studies (54), though long-term deleterious consequences, particularly in the setting of heart disease, remain unexplored.

The increased protein synthesis observed in the heart and loss of lean muscle mass in the AngII+Lira mice may suggest that circulating amino acids are being preferentially taken up by the heart or scavenged from peripheral muscle. We performed plasma metabolomic studies in all 3 groups and found that the AngII+Lira mice had elevated circulating amino acids (6 of 23 metabolites elevated in AngII+Lira vs sham mice, Supplemental Figures 5 and 6). Regardless of the source of the elevated circulating amino acids, these findings raise concerns about GLP-1 receptor agonists related to sarcopenia and frailty, particularly given the high prevalence of diastolic dysfunction in the community (55) and the growing use of GLP-1 in weight-loss therapies (56).

The metabolomic screening of cardiac tissue implicated less pronounced changes to other metabolites. Though glycolytic and citric acid cycle intermediates were largely unchanged between the treatment groups, one notable finding included a significant accumulation of Acetyl-CoA in the AngII cohort, which was normalized in the AngII+Lira mice. Acetyl-CoA accumulation may lead to feedback inhibition of FAO (57), which has previously been reported in a similar model of diastolic dysfunction (22). Despite restoring Acetyl-CoA in the AngII+Lira cohort, we could not detect any correction of FAO in cardiac homogenates with labeled palmitate (Figure 5). In fact, the AngII+Lira cohort had significantly lower FAO than the sham cohort, and lower palmitate oxidation to CO_2_ compared to the AngII mice. It remains possible that acetyl-CoA levels normalize in AngII+Lira by reducing beta-oxidation or through the consumption of Acetyl-CoA in the TCA cycle.

In summary, the study presented herein demonstrates that Ang-treated mice develop diastolic disease through multiple echocardiographic parameters, and that daily liraglutide treatment completely prevents these markers of diastolic dysfunction. These changes occur independently of blood pressure or hypertrophy but are associated with lower myocardial fibrosis. Metabolomic changes found dramatic changes to amino acid accumulation in the hearts of AngII+Lira mice, which may promote protein synthesis to support cardiac remodeling against afterload in the presence of angiotensin. However, the liraglutide treatment was also associated with losing lean muscle mass, which may be deleterious in the long term. Future studies are needed to explore the long-term consequences of GLP-targeted therapies, including the risk of sarcopenia and frailty. Additional studies examining the bioenergetic effects of amino acid build-up, and possibly anaplerotic roles of these metabolites, in this model.

## Abbreviations

4EBP1: Eukaryotic translation initiation factor 4E-binding protein 1
AngII: Angiotensin II
ANOVA: Analysis of variance
ASM: Acid soluble metabolites
BCAA: Branched-chain amino acids
BCKDH: Branched-chain α-keto acid dehydrogenase complex
BP: Blood pressure
CPT1b: Carnitine palmitoyltransferase 1B
ECG: Electrocardiogram
EF: Ejection Fraction
ET: Ejection Time
FAO: Fatty acid oxidation
GAPDH: Glyceraldehyde 3-Phosphate Dehydrogenase
GLP-1: Glucagon-like peptide 1
Gluc 6p: Glucose 6 phosphate
Gly 3p: Glyceraldehyde-3-phosphate
HFpEF: Heart failure with preserved ejection fraction
H&E: Hematoxylin and eosin
IP: Intraperitoneal
IVCT: Isovolumic contraction time
IVRT: Isovolumic relaxation time
KCL: Potassium Chloride
LC-HRMS: Liquid chromatography-high resolution mass spectrometry
Lira: liraglutide
LV: Left Ventricle
MCAD: Medium chain acyl-CoA dehydrogenase
P62: p62/Sequestosome 1
Rib 5p: Ribulose-5-phosphate
rLSR: Reverse longitudinal strain rate
S6: S6 Ribosomal Protein
Sed 7p: Seduheptulose 7-phosphate
VLCAD: Very long chain acyl-CoA dehydrogenase

## Acknowledgements

CR, MJ, EG, and BK were responsible for conceptualizing these studies. CR designed the methodology and with KR, ML, and SM, performed the investigation. CR, SM, SW, MJ, and EG were responsible for data curation. Formal analyses were performed by CR, SM, SW, MJ, EG, and BK. Visualization was completed by CR, SM, MJ, and EG. CR and BK wrote the manuscript. BK supervised the project and provided resources for its completion. All authors reviewed the final manuscript and are responsible for its integrity.

## Sources of Funding

Research reported in this manuscript was supported by: American Heart Association Transformational Project Award 18TPA34230048, NIH Instrument Grant for Advanced High-Resolution Rodent Ultrasound Imaging System 1S10OD023684-01A1, Veteran’s Administration Grant IK2BX005785, and NIH NRSA 1F32HL156428.

## Disclosures

None of the authors have financial disclosures or conflicts of interest.

## Notes

### Competing Interest Statement

The authors have declared no competing interest.

### Summary of Updates

Updated Figures 2 to include additional measures of hypertrophy, rearranged Figure 3 and supplemental Figures 2-4, and added text to manuscript to reflect clinical relevance.

