## Supplemental Figures for "Liraglutide protects against diastolic dysfunction and improves ventricular protein translation"

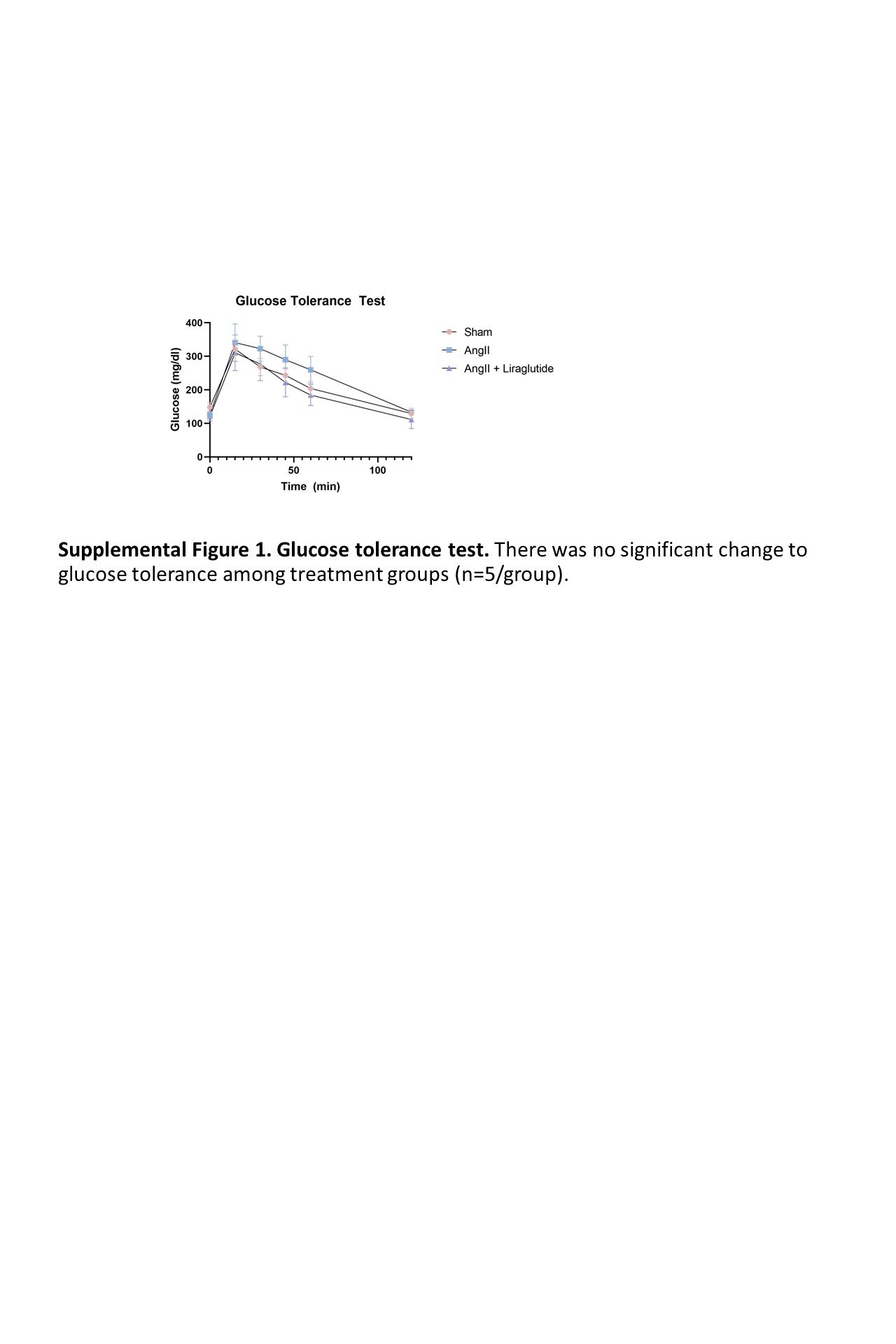

**Supplemental Figure 1. Glucose tolerance test.** There was no significant change to glucose tolerance among treatment groups (n=5/group).

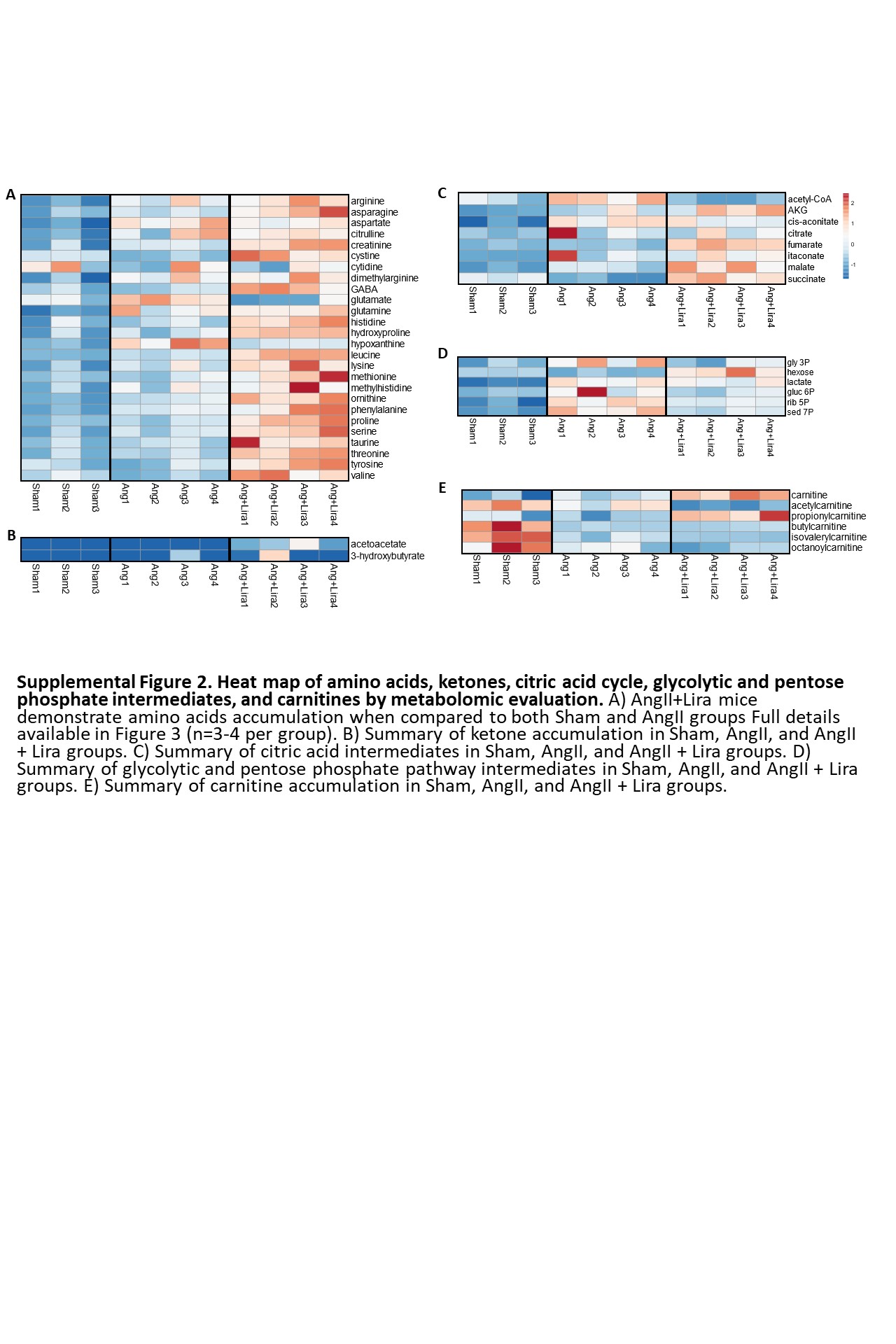

**Supplemental Figure 2. Heat map of amino acids, ketones, citric acid cycle, glycolytic and pentose phosphate intermediates, and carnitines by metabolomic evaluation.** A) AngII+Lira mice demonstrate amino acids accumulation when compared to both Sham and AngII groups Full details available in Figure 3 (n=3-4 per group). B) Summary of ketone accumulation in Sham, AngII, and AngII + Lira groups. C) Summary of citric acid intermediates in Sham, AngII, and AngII + Lira groups. D) Summary of glycolytic and pentose phosphate pathway intermediates in Sham, AngII, and AngII + Lira groups. E) Summary of carnitine accumulation in Sham, AngII, and AngII + Lira groups.

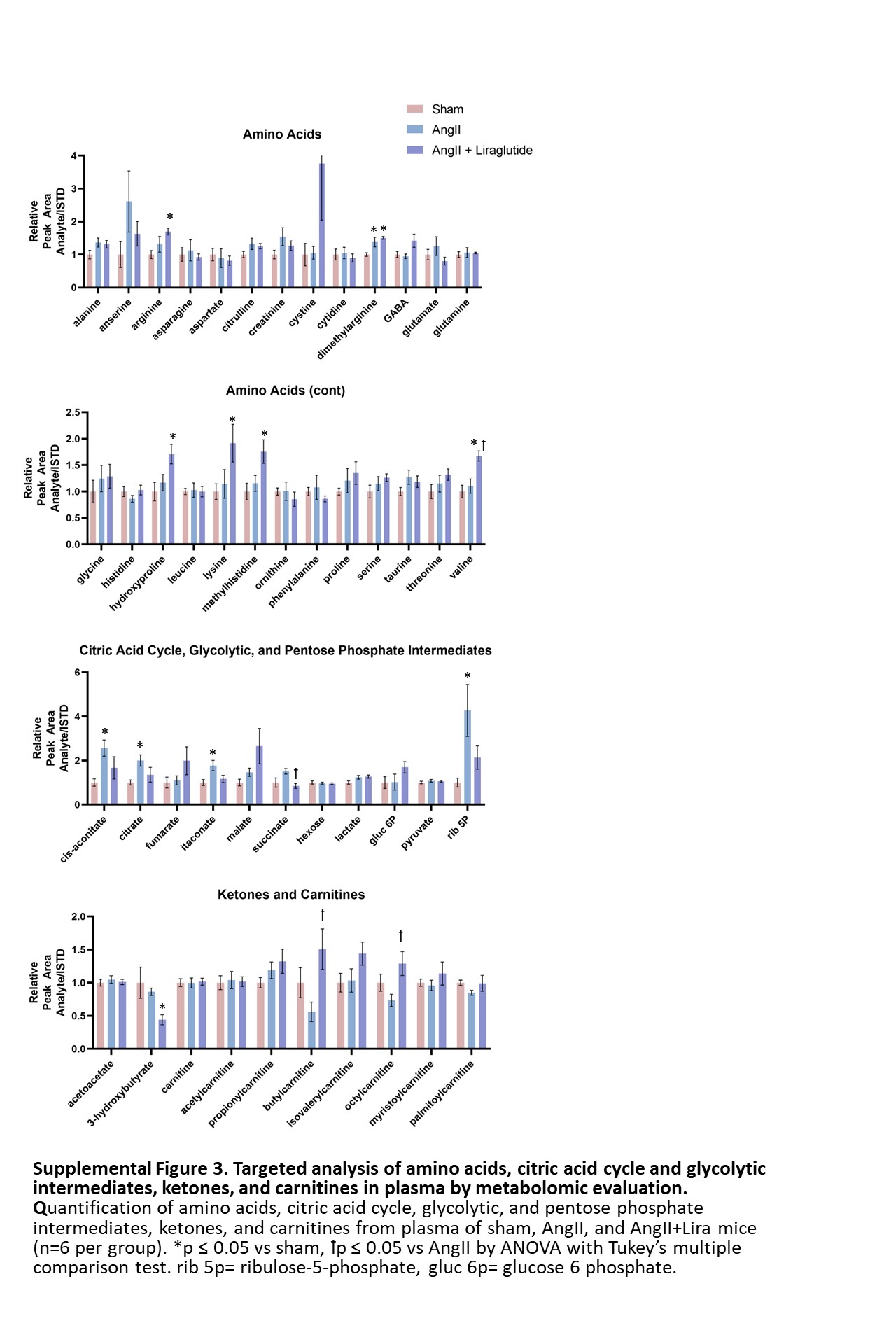

**Supplemental Figure 3. Targeted analysis of amino acids, citric acid cycle and glycolytic intermediates, ketones, and carnitines in plasma by metabolomic evaluation.** Quantification of amino acids, citric acid cycle, glycolytic, and pentose phosphate intermediates, ketones, and carnitines from plasma of sham, AngII, and AngII+Lira mice (n=6 per group). *p ≤ 0.05 vs sham, ꝉp ≤ 0.05 vs AngII by ANOVA with Tukey’s multiple comparison test. rib 5p= ribulose-5-phosphate, gluc 6p= glucose 6 phosphate.

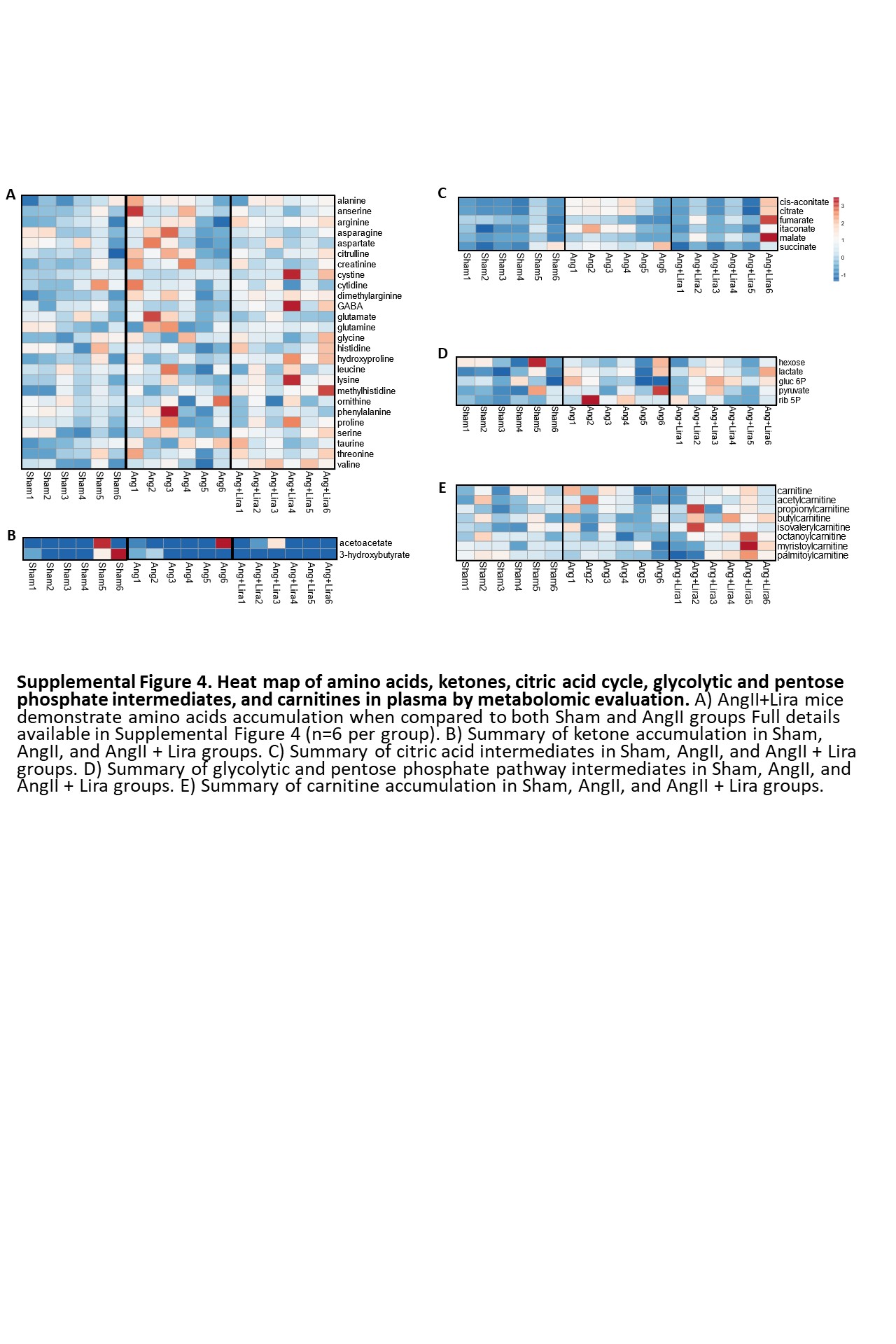

**Supplemental Figure 4. Heat map of amino acids, ketones, citric acid cycle, glycolytic and pentose phosphate intermediates, and carnitines in plasma by metabolomic evaluation.** A) AngII+Lira mice demonstrate amino acids accumulation when compared to both Sham and AngII groups Full details available in Supplemental Figure 4 (n=6 per group). B) Summary of ketone accumulation in Sham, AngII, and AngII + Lira groups. C) Summary of citric acid intermediates in Sham, AngII, and AngII + Lira groups. D) Summary of glycolytic and pentose phosphate pathway intermediates in Sham, AngII, and AngII + Lira groups. E) Summary of carnitine accumulation in Sham, AngII, and AngII + Lira groups.

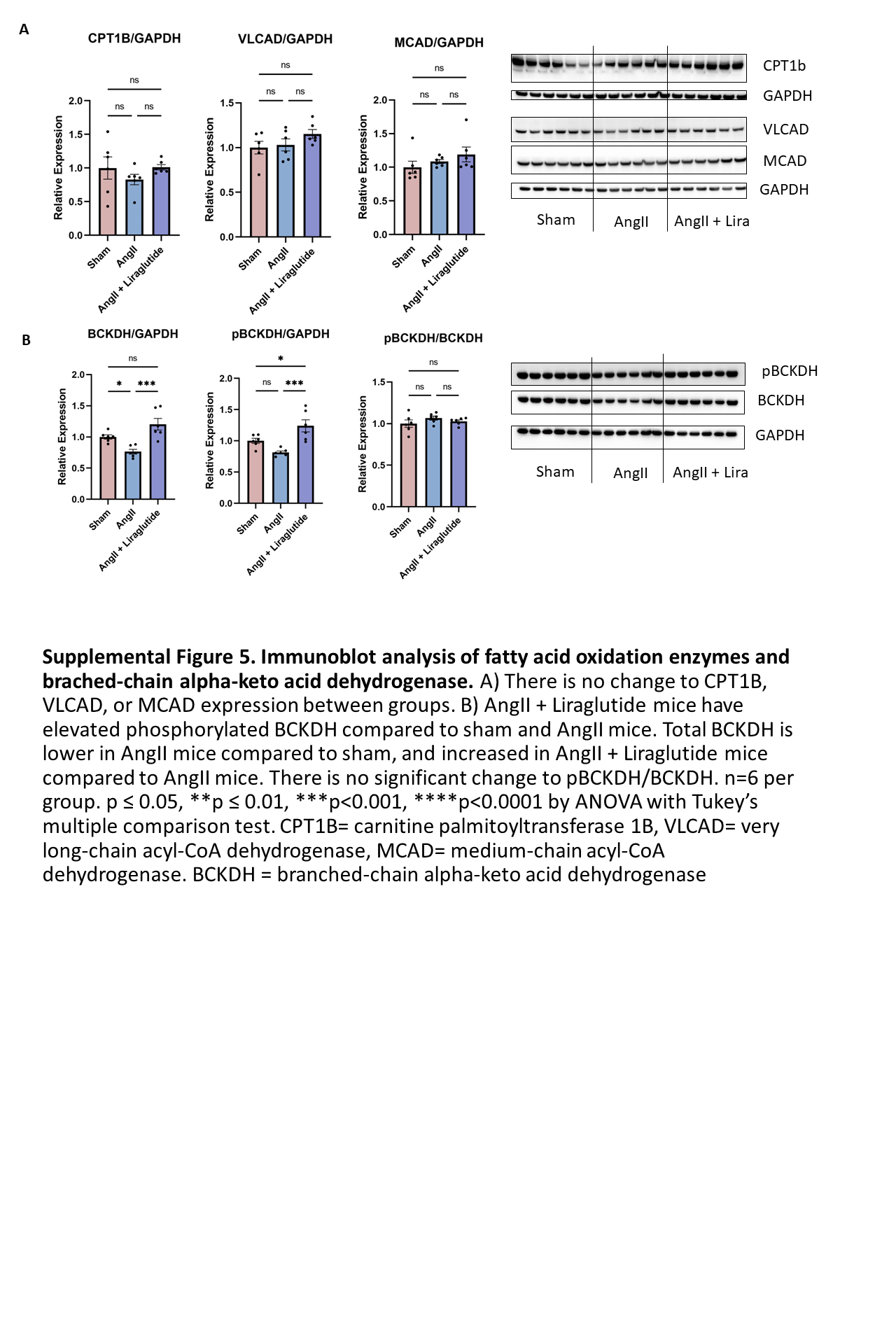

**Supplemental Figure 5. Immunoblot analysis of fatty acid oxidation enzymes and brached-chain alpha-keto acid dehydrogenase.** A) There is no change to CPT1B, VLCAD, or MCAD expression between groups. B) AngII + Liraglutide mice have elevated phosphorylated BCKDH compared to sham and AngII mice. Total BCKDH is lower in AngII mice compared to sham, and increased in AngII + Liraglutide mice compared to AngII mice. There is no significant change to pBCKDH/BCKDH. n=6 per group. p ≤ 0.05, **p ≤ 0.01, ***p<0.001, ****p<0.0001 by ANOVA with Tukey’s multiple comparison test. CPT1B= carnitine palmitoyltransferase 1B, VLCAD= very long-chain acyl-CoA dehydrogenase, MCAD= medium-chain acyl-CoA dehydrogenase. BCKDH = branched-chain alpha-keto acid dehydrogenase.

**Supplemental Figure 6. Original and uncut blots.** A) Western blots from Figure 4. B) Western blots from Supplemental Figure 5.

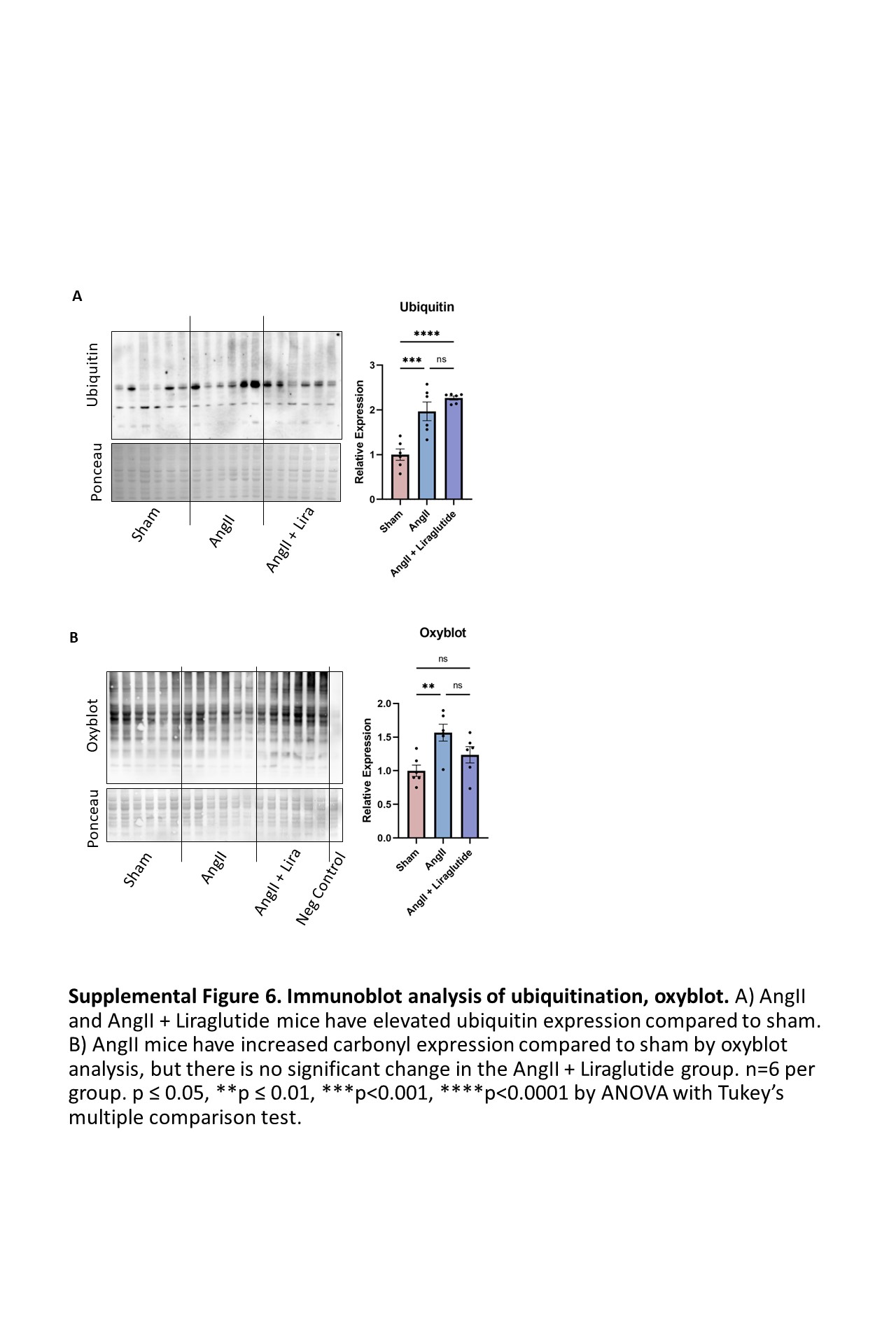

**Supplemental Figure 6. Immunoblot analysis of ubiquitination, oxyblot.** A) AngII and AngII + Liraglutide mice have elevated ubiquitin expression compared to sham. B) AngII mice have increased carbonyl expression compared to sham by oxyblot analysis, but there is no significant change in the AngII + Liraglutide group. n=6 per group. p ≤ 0.05, **p ≤ 0.01, ***p<0.001, ****p<0.0001 by ANOVA with Tukey’s multiple comparison test.
